# Advanced meditation stabilizes cortical dynamics through inhibitory gain control linked to absorptive depth

**DOI:** 10.64898/2026.11.08.765124

**Authors:** Rajanikant Panda, Gulshan Kumar, Pablo Núñez, Balachundhar Subramaniam, Kalpana Bhandare, Saketh Malipeddi, Genevieve Major, Olivia Gosseries, Steven Laureys, Prejaas Tewarie

## Abstract

Advanced meditation can shift conscious experience from ordinary body- and world-centered awareness toward absorption, detachment and dissociation-like states, yet the neural mechanisms that stabilize these immersive states remain unclear. We studied 18 long-term meditators during eyes-open rest and Rajyoga meditation, a practice in which practitioners reduce bodily identification while enhancing inner self-awareness. We combined multidimensional neural approach including frontal EEG spectral power, temporal non-stationarity and corticothalamic neural-field modeling, because these measures probe underlying complementary levels of the brain state transition: the strength of neural activity, its temporal stability, and the circuit level inhibition-excitation gain mechanisms. This framework is especially relevant to meditation because oscillatory, temporal organization may distinguish stable meditative states, and corticothalamic dynamics are closely linked to arousal and conscious state regulation. We correlated meditation induced neural changes with subjective ratings of absorption and dissociation. Compared with rest, meditation was associated with higher beta1, beta2 and gamma power, with no change in the slower bands (delta, theta and alpha), and with lower beta-band temporal non-stationarity, indicating a more stable expression of fast cortical dynamics. Fitting the corticothalamic model to the EEG spectra showed stronger cortical inhibitory gain and lower effective intracortical gain, placing meditation in an inhibition-dominant regime, with lower gain in the loop between excitatory and inhibitory neurons within the cortex. During meditation, these model estimates were related to subjective absorption and showed trend level links with dissociation. Together, these findings suggest that Rajyoga meditation stabilizes cortical dynamics through inhibitory gain control, linking absorption (deep and steady attention on the meditation) to dissociation-like self-detachment.

## 1. Introduction

Meditation offers a rare experimental window into how the brain can voluntarily reshape conscious experience (Braboszcz et al., 2017; Saketh et al., 2026). With sustained practice, attention can become more stable, spontaneous self-referential mentation can recede, and ordinary body- and world-centered awareness may give way to states marked by deep absorption, self-detachment, and altered modes of presence. Although such experiences are central to contemplative traditions, their neurophysiological basis remains only partly understood. Electroencephalography is especially well suited to tackle this problem because it captures ongoing neural dynamics at the timescale on which meditative states emerge, stabilise, and fluctuate. Over the past two decades, EEG studies have helped shift meditation research away from purely descriptive phenomenology toward a more mechanistic account of how trained mental practice reorganises cortical activity in real time.

Rajyoga meditation, as taught in the Brahma Kumaris tradition, provides a particularly informative model for studying how the brain supports deep, inwardly focused meditative states. Unlike practices centered primarily on breath monitoring, interoceptive tracking, or sensory withdrawal, Rajyoga is typically practiced with the eyes open and involves a deliberate shift of attention away from identification with the physical body and external environment toward an inwardly sustained awareness of the self as peaceful and consciously present, a state referred in Rajyoga practice as “soul consciousness’’. (Kumar et al., 2023; Panda et al., 2016; Singh et al., 2026). Rajyoga therefore brings together two experiential dimensions usually studied separately: absorption and dissociation-like detachment. Absorption refers to deep attentional immersion in the meditative state, whereas dissociation-like detachment, used here in a non-pathological sense and close to what is also called self-detachment or de-identification, refers to a loosening of ordinary body- and world-anchored self-experience (Kumar et al., 2023). Rajyoga aims to stabilize awareness while reducing habitual identification with bodily sensations, external cues, and self-referential thought (Babu et al., 2020; Kumar et al., 2023; Panda et al., 2016; Singh et al., 2026). It therefore offers a precise model for testing how absorptive self-detachment is implemented in cortical dynamics.

Neurophysiology and imaging studies on Rajyoga meditation suggest that this practice engages brain systems relevant to both self-processing and cognitive control. In a combined EEG-fMRI study, Panda and colleagues reported alterations in the temporal dynamics of the default mode network, a network strongly linked to mind-wandering, autobiographical mentation, and self-referential thought (Panda et al., 2016). Long-term practitioners also differed from non-meditators in grey matter volume across the frontal, insular, and precuneal regions, as measured with structural MRI (voxel-based morphometry). Practitioners had more grey matter in the right superior frontal gyrus, left orbitofrontal cortex and precuneus, and their happiness scores were linked to grey matter volume in frontal, orbitofrontal, insular and anterior cingulate regions (Babu et al., 2020). More recently, EEG network theory work has suggested altered information exchange within and across hemispheres during eyes-open Rajyoga meditation (Kumar et al., 2023). These studies show that Rajyoga engages self-referential and attentional networks. However, the cortical mechanism behind these changes is still unknown. For example, it is not clear whether meditation increases excitation in the cortex or changes the balance between excitation and inhibition.

A common EEG finding in meditation research is that higher-frequency activity, especially in the gamma band, changes during practice. Long-term practitioners in multiple traditions show elevated gamma-band power, both during practice and as a resting trait, and higher gamma power has been associated with more meditation experience (Braboszcz et al., 2017; Ferrarelli et al., 2013; Lutz et al., 2004). However, the findings are not fully consistent, and lower gamma power has also been reported in advanced meditators. Gamma activity is also linked to focused and sustained attention (Jensen et al., 2007a; Ray et al., 2008). Because absorption involves deep and steady attention, gamma activity may also be linked to absorptive states. However, to our knowledge, no study has yet tested this link directly, and this is one aim of the present study. We therefore expected Rajyoga meditation to raise power in the higher-frequency bands relative to rest. Temporally averaged band power captures only how much energy is in a frequency band, but not how steady that energy. A state built on continuous, undistracted attention should increase the temporal regularity of the signal as much as its high-frequency power. Work that has approached meditation through complexity and entropy points in this direction, reporting more orderly and less variable EEG dynamics in experienced practitioners (Huang & Lo, 2009; Walter & Hinterberger, 2022) and showing that skilled meditators can shift these dynamics rapidly and at will (Nair et al., 2017). We captured this property with a Kullback–Leibler divergence measure of non-stationarity, which asks how far the moment-to-moment distribution of spectral energy departs from an even one across the epoch (Bhattacharya & Petsche, 2001; Núñez et al., 2017, 2019; Tong et al., 2007). If Rajyoga meditation reflects a more stable attentional regime, it should reduce this non-stationarity and do so most clearly in the bands that support sustained engagement.

Power and non-stationarity are purely descriptive measures of an electroendephalographic signal, but they do not provide any information on their specific neural generators. To obtain estimators of the underlying generators behind the observed signals, we used the corticothalamic model developed by Robinson and colleagues. This model can reproduce realistic EEG spectra using a small number of physiological parameters. It describes four connected groups of neurons, two in the cortex and two in the thalamus. Among them are the gains of the cortical excitatory and inhibitory populations, the gains of the corticothalamic loops, and a normalized intracortical gain that summarizes the balance between them. The model parameters have been observed to shift as brain state changes, for instance with arousal (Abeysuriya & Robinson, 2016; Robinson et al., 1997, 2001, 2002; Sanz-Leon et al., 2018; Van Albada & Robinson, 2013), as well as in altered states of consciousness, including disorders of consciousness (Assadzadeh et al., 2023; Cóngyŭ et al., 2025; Tewarie et al., 2025). Fitting the model to the rest and meditation spectra allows us to describe the meditative shift in terms of estimated physiological quantities, such as cortical inhibition and gain. Gamma oscillations are produced by the interaction between excitatory and inhibitory neurons, with inhibitory neurons setting their rhythm (Buzsáki & Wang, 2012). Stronger cortical inhibition has also been proposed as a source of the higher gamma power seen in meditators (Biswas et al., 2026). We therefore expected meditation to shift the cortex toward stronger inhibition. In the model, this would appear as a more negative cortical inhibitory gain, G_ei_, which is the strength of the input from inhibitory to excitatory cortical neurons. We also expected a lower value of X = G_ee_/(1 − G_ei_), where G_ee_ is the gain of the excitatory connections within the cortex. X is not a separate parameter of the model; it is a summary measure computed from the two cortical gains, which shows how strongly excitation within the cortex is offset by inhibition (Robinson et al., 2002).

While physiological measurements and estimations offer a descriptive view of neuroimaging changes, it is important to connect these shifts to what practitioners subjectively report feeling. Absorption and dissociation are two experiences often reported in deep meditation, as described above (Panda et al., 2023; Rousseaux et al., 2023) (Vago & David, 2012). The two are closely related (Bowins, 2012; Soffer-Dudek et al., 2015), and deeper absorption has been proposed to lead to deeper altered states of consciousness (Mohr, 2018). Inhibitory neurons help the cortex select which signals it responds to, and they support the gamma rhythm that is linked to attention (Cardin et al., 2009; Sohal et al., 2009). Stronger inhibition could therefore help to keep attention fixed on meditation and reduce the effect of bodily and outside signals. Based on this, we expected that participants who are more deeply absorbed during meditation should show a stronger inhibitory shift, and those whose experience changes mostly from rest to meditation should show the largest change in the parameters. This approach, which links brain measures to first-person reports, follows the neurophenomenology framework (Lutz & Thompson, 2003; Varela, 1996).

In the present study, we recorded frontal EEG from experienced Rajyoga meditators during eyes-open rest and soul-conscious meditation state to test a mechanistic account of this state. We measured three features of the EEG: spectral power, temporal non-stationarity, and the balance between excitation and inhibition estimated with the corticothalamic model. Together, these measures show how much fast activity there is, how stable it is over time, and which circuit changes may produce it. Based on the studies reviewed above, we expected that meditation would (1) increase fast (beta and gamma) activity, (2) make this activity more stable over time, and (3) shift the cortex toward stronger inhibition. We also tested whether these changes were related to how absorbed and detached participants felt during meditation.

## 2. Methods

### 2.1 Participants and experimental design

Eighteen experienced Raja Yoga meditators from the Brahma Kumaris tradition participated in the study (11 men, 7 women; mean age = 48.7 years, SD = 8.3, range = 31 to 55). Participants were long-term practitioners with an average of 23.9 years of meditation experience (SD = 8.0) and 2.2 hours of daily practice (SD = 1.1, range 0.5-4.0). Estimated lifetime practice, calculated as years of practice × daily hours × 365, was a mean of about 20,990 hours (SD 14,320, range 2,190 to 44,710). Mean formal education was 16.6 years (SD 2.5, range 14-23). The data were acquired at Global Hospital and Research Center, Mount Abu, Rajasthan, India. The Inclusion criteria was participants should be age between 25 to 60 years, minimum schooling education (12 year education), advanced meditation practice with every day at list one hours, and at list 10 years of practice, no history of neurological or psychiatric illness, while the exclusion criteria was: if the participants have minimum 3-month of discontinuity in meditation practice in last three years, any neuropsychiatric disorders, or practice any other kind of meditation/physical activity extensively. The study was approved by the Global Hospital and Research Center Ethics Committee. All participants gave informed consent, and all procedures were conducted in accordance with the Declaration of Helsinki.

In this study, advanced meditation refers to extensive contemplative experience and sustained engagement in practices aimed at cultivating deep meditative absorption and self-detachment. Importantly, participants were not full-time meditation center residents but community dwelling practitioners who maintained their usual family, social, and professional responsibilities while incorporating prolonged daily meditation into their lives. In addition to their regular practice, participants undergo intensive meditation retreats lasting at least 15 consecutive days annually. During these retreats, they substantially reduced external engagement, including social media use, verbal communication, and other everyday interactions, to facilitate sustained inward attention, deeper absorption, and dissociation-like detachment from the external environment. Thus, the cohort represented experienced meditators whose contemplative training combined long-term daily practice with periodic intensive retreats.

Participants underwent continuous EEG recording during two sequential conditions. First, a 5-minute eyes-open resting-state baseline was acquired, during which participants were instructed to stay awake, relaxed, and still, without engaging in any specific meditative practice. This was followed by a 5-minute eyes-open soul-conscious meditation condition. Soul-conscious meditation is a core practice within the Rajyoga tradition, in which practitioners intentionally reduce identification with the physical body and the external sensory environment while cultivating inner self-awareness (Panda et al., 2016; Kumar et al., 2023; Singh et al., 2026). Practitioners often conceptualize this state as awareness of the self as a point of light associated with peace, purity, and positive virtues. This design yielded a paired, within-subject comparison between resting baseline and meditation, allowing each participant to serve as their own control. An overview of the recording and analysis pipeline is shown in Figure 1.

**Figure 1.**
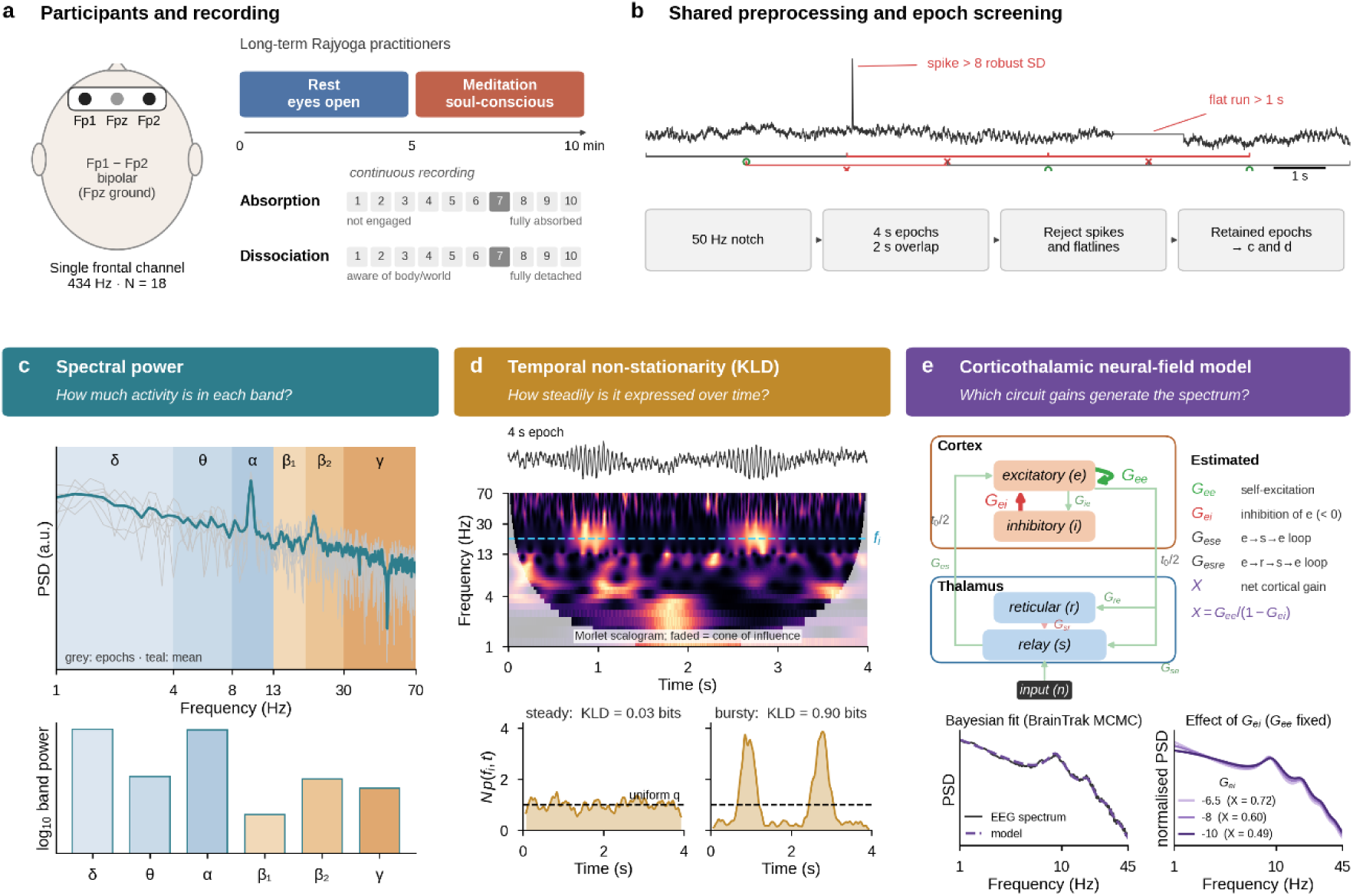
Analysis pipeline. A) Frontal single-channel EEG (Fp1–Fp2) recorded during eyes-open rest and Rajyoga meditation (5 min each; N = 18), followed by absorption and dissociation ratings. B) Preprocessing: notch filtering, 4 s epochs and artifact rejection. C) Spectral power: epoch-averaged periodograms integrated within six frequency bands. D) Temporal non-stationarity: Kullback–Leibler (KL) divergence between the temporal energy distribution of the wavelet scalogram and a uniform distribution. Near zero values mean steady activity, while higher values mean bursty activity. E) Corticothalamic neural-field model fitted to each spectrum to estimate cortical gains and, loop gains and, and net intracortical gain. Signals are simulated for illustrative purposes.

### 2.2 EEG acquisition

EEG was recorded with the Neurosteer single-channel system (Neurosteer Inc., New York, NY), a portable frontal EEG device that has been validated for extracting clinically meaningful features from a single channel (Molcho et al., 2023). An FDA-cleared adhesive patch carrying three electrodes was placed on the participant’s forehead with a conductive dry gel. The electrodes were positioned over the prefrontal region according to the international 10/20 system, and a single bipolar EEG channel was derived from the difference between Fp1 and Fp2, with a ground electrode positioned at Fpz. Signals were digitized at 434 Hz. Electrode contact quality was checked before recording, and participants were seated comfortably and instructed to remain still and minimize blinking, jaw movement, and speech. Recording was continuous across both conditions, yielding 10 minutes of data per participant.

### 2.3 Subjective behavioral measures

Two subjective experiential dimensions were assessed: dissociation and absorption. Dissociation refers to a temporary alteration or partial decoupling of normally integrated mental processes, including awareness, memory, emotion, bodily experience, and sense of agency. Absorption refers to the depth of attentional engagement in an internal experience, including focused awareness, imagery, meditative content, and contemplative self-reflection. In this study, it was treated as a subjective index of meditative depth. Both were measured with simple single-item ratings.

Immediately after each condition (that is, once after the rest period and once after the meditation period), participants rated the depth of absorption and the degree of dissociation they had experienced during that condition on a 1-to-10 numerical rating scale. For dissociation, 1 indicated full awareness of the body and external environment and 10 indicated complete subjective detachment from bodily sensations and environmental stimuli. For absorption, 1 indicated no engagement with meditative or internally focused content, and 10 indicated complete absorption in the meditative state or associated internal experience. Participants also reported the approximate time taken to reach the rest and meditative states. This yielded absorption and dissociation scores for both conditions, together with a rest-to-meditation change score for each measure.

### 2.4 Preprocessing and epoch screening

Preprocessing was kept to a minimum, because only one EEG channel was available, and it was applied identically across all analyses, so that every measure was derived from the same retained data. A 50 Hz zero-phase IIR notch filter (quality factor 35) was applied to attenuate line noise. No other filter was applied, and the data were not resampled (sampling rate 434 Hz). Each five-minute condition was then divided into 4-second epochs with 2-second overlaps. No independent component analysis or component-based artifact classification was used, because these methods need several channels to separate brain and non-brain sources, and our recording had only one channel. Instead, epochs with spikes or flat segments were removed (see below). Muscle activity can still add power at high frequencies, mainly in the gamma band, and we discuss this in the limitations.

Epochs were screened for two artifact types on the raw signal, before the notch filter was applied. Transient spikes were detected with a robust z-score based on the median and the median absolute deviation, and an epoch was rejected if any sample exceeded eight robust standard deviations from the epoch median. Flatline segments were detected as runs of near-constant signal, and an epoch was rejected if its longest flat run exceeded one second. All remaining epochs were retained. Because rejection was performed once per participant and condition, the spectral-power and non-stationarity analyses were computed from exactly the same epochs. We verified that the total, kept, rejected, spike, and flatline epoch counts matched across the two pipelines. Epoch retention was high, with a median rejection rate below 2% per condition.

### 2.5 Spectral power

For each retained epoch, the power spectral density was estimated with a periodogram (Hann window, constant detrend, density scaling), and the epoch spectra were averaged within each participant and condition. Band power was obtained by integrating the averaged spectrum over six frequency bands: delta (1-4 Hz), theta (4-8 Hz), alpha (8-13 Hz), beta1 (13-19 Hz), beta2 (19-30 Hz), and gamma (30-70 Hz). Band-power values were log10-transformed before group-level statistics.

### 2.6 Temporal non-stationarity

Temporal non-stationarity was quantified with the Kullback-Leibler divergence (KLD) between the CWT and a uniform distribution, computed on the same artifact-screened epochs, following the approach first described by Tong et al. (2007) and later used by Núñez et al. (2017). A signal is weakly stationary when its spectral content does not change over time. This implementation of KLD captures departures from this condition by measuring how unevenly energy is distributed across the epoch at each frequency.

Each 4 s epoch was decomposed with a continuous wavelet transform using a complex Morlet wavelet (cmor1-1), producing a time-frequency representation (TFR, or wavelet scalogram) over 139 pseudo-frequencies spanning 1–70 Hz in 0.5 Hz steps, with scales derived from the wavelet’s center frequency and the 434 Hz sampling rate. A cone of influence mask was applied to discard edge-affected coefficients, so that at each frequency only the valid time points contributed to the calculation (Torrence & Compo, 1998).

The marginal frequency distribution and the time-frequency representation (TFR) are described in Eqs. (1) and (2), respectively, while the KLD is described in Eq. (3)

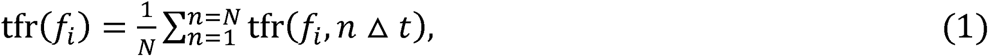

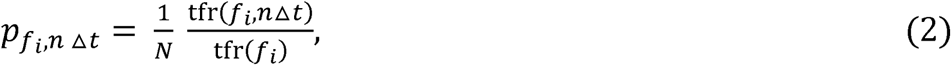

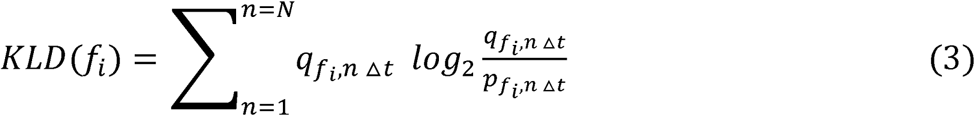

In these expressions, tfr(*f_i_*) is the value of the time–frequency representation at frequency fi and time point n, here the wavelet scalogram, and N is the number of time points that contribute at that frequency after cone of influence masking. The reference distribution 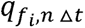 is uniform in time, so Eq. (3) quantifies how far the observed distribution of energy departs from an even spread across the epoch. A stationary signal, whose energy is spread evenly in time, yields a temporal distribution close to uniform and a KLD near zero, whereas energy that is concentrated at particular moments produces larger values (Núñez et al., 2017). The per-frequency KLD values were averaged within each of the six frequency bands (delta, theta, alpha, beta1, beta2, gamma) for every epoch and then averaged across the retained epochs to give one value per participant, condition, and band.

### 2.7 Corticothalamic neural-field modeling

We fitted the corticothalamic neural-field model of Robinson and colleagues (Robinson et al., 1997, 2001, 2002) to each participant’s rest and meditation power spectra. The model represents the brain as four interacting neural populations: cortical excitatory (*e*) and cortical inhibitory (*i*) neurons, thalamic relay neurons (*s*), and thalamic reticular neurons (*r*), together with an external input (*n*). Each population converts incoming activity into a mean firing rate through a sigmoidal response function, and signals pass between populations subject to synaptodendritic filtering, corticothalamic conduction delays, and cortical wave propagation. From this compact set of physiological ingredients, the model predicts the EEG power spectrum in closed form. A full mathematical derivation of the model as well as its steady-state frequency representation is detailed in (Robinson et al., 2004; Abeysuriya et al., 2014).

The shape of the modeled spectrum is governed by a small set of gain parameters. The gain *G_ab_* represents the response produced in population a per unit input from population b, where a and b denote any two of the model’s populations, and connections mediated by inhibitory neurons take negative values. We analyzed the cortical excitatory gain (*G_ee_*) and the cortical inhibitory gain (*G_ei_*), together with the loop gains of the corticothalamic system: the excitatory corticothalamic loop gain (*G_ese_*, cortex → relay → cortex) and the inhibitory corticothalamic loop gain (*G_esre_*, cortex → reticular → relay → cortex). Because *G_ei_* is the gain of the inhibitory-to-excitatory cortical connection, it is negative by construction, so a more negative value denotes stronger cortical inhibition. To summarize the cortical dynamics in a single quantity, we used the normalized parameter *X= G_ee_/(1-G_ei_)*, which corresponds to the cortical (intracortical) loop gain of the model (Robinson et al., 2002). X is not a separate parameter of the model; it is computed from G_ee_ and G_ei_. A lower value indicates a smaller net intracortical gain, that is, a cortical loop in which excitation is more strongly offset by inhibition.

For the modeling analysis, power spectra were estimated separately from the specific preprocessing used by the fitting procedure. The single-channel signal for each condition was divided into 60-second blocks, and the power spectral density of each block was computed using Welch’s method (4-second Hamming windows, 1-second overlap, FFT length equal to 4 times the sampling rate) and lightly smoothed with a 10-point moving average. As in the spectral-power and non-stationarity analyses, no band-pass filter was applied before spectral estimation; instead, the spectra were retained over the range required by the model fit, approximately 0.75–47 Hz. This yielded one power spectrum per 60-second block for each participant and condition.

Model parameters were estimated from these spectra using BrainTrak, an automated Bayesian model-inversion procedure (Abeysuriya & Robinson, 2016). BrainTrak treats spectral fitting as a problem of statistical inference, using a Metropolis–Hastings Markov-chain Monte Carlo scheme to draw samples from the posterior distribution of the parameters given the observed spectrum, over an effective fit range of approximately 1–45 Hz. Fit quality is scored by a frequency-weighted fractional difference between the modeled and empirical spectra (weighted so that each frequency decade contributes equally), The posterior is set to zero for physiologically implausible parameter combinations or that place the system beyond its stability boundary, so that every accepted fit corresponds to a stable, physiologically admissible brain state. Because BrainTrak fits the successive blocks as a tracked sequence, using each fit to initialize the next, the first three blocks were treated as burn-in and discarded; the parameter estimates from the remaining per-block fits were then averaged to give a single value per participant and condition. Across participants, the fitted model reproduced the observed spectra closely, with goodness-of-fit values in the range χ² ≈ 1–2 (Supplementary Figure S1).

## 3. Statistical analysis

All paired rest-versus-meditation contrasts used the two-sided Wilcoxon signed-rank test. This choice was made because Shapiro-Wilk tests indicated that several paired-difference distributions departed from normality, and because the sample size was modest (N=18). The non-parametric test was therefore applied consistently across all measures. For each contrast, we report the median in each condition, the median paired difference (meditation minus rest), a 95% confidence interval for the median difference obtained from 3,000 bootstrap resamples, the Wilcoxon statistic (W), an approximate effect size r, and the p-value. Within each analysis family, that is, the six spectral bands for the relative power, the six KLD bands, and the five model parameters, p-values were corrected for multiple comparisons using the Benjamini-Hochberg false-discovery-rate (FDR) procedure, and effects were considered significant at an FDR-corrected threshold of 0.05.

Brain-behavior relationships were assessed with Spearman rank correlations, with 95% confidence intervals for each correlation obtained from 3,000 bootstrap resamples (N = 18). We tested the two model parameters of main interest, *G_ei_* and *X*, for correlation with absorption and dissociation, both in the meditation state and as rest-to-meditation change scores.

Spectral power, statistical, and correlation analyses were carried out in Python (NumPy, SciPy), and wavelet-based non-stationarity analysis and neural field model fitting were carried out in MATLAB (version R2022a; The MathWorks, Natick, MA). Model fitting used the BrainTrak toolbox (https://github.com/BrainDynamicsUSYD/braintrak)

## 4. Results

### 4.1 Meditation increases beta and gamma spectral power

Meditation was associated with a selective increase in spectral power in the higher-frequency bands, with the magnitude of the effect increasing progressively from beta to gamma frequencies (Figure 2). After FDR correction across the six bands, beta1, beta2, and gamma power were all higher during meditation than at rest, each with a large effect size: beta1 increased by 0.088 log10 units (95% CI [0.019, 0.238]; W = 27, r = 0.60, p = 0.009, FDR p = 0.018), beta2 by 0.197 ([0.060, 0.378]; W = 19, r = 0.68, p = 0.002, FDR p = 0.007), and gamma by 0.262 ([0.090, 0.391]; W = 15, r = 0.72, p = 0.001, FDR p = 0.006). The three lower bands did not differ between conditions (delta, theta, and alpha all FDR p > 0.28).

**Figure 2.**
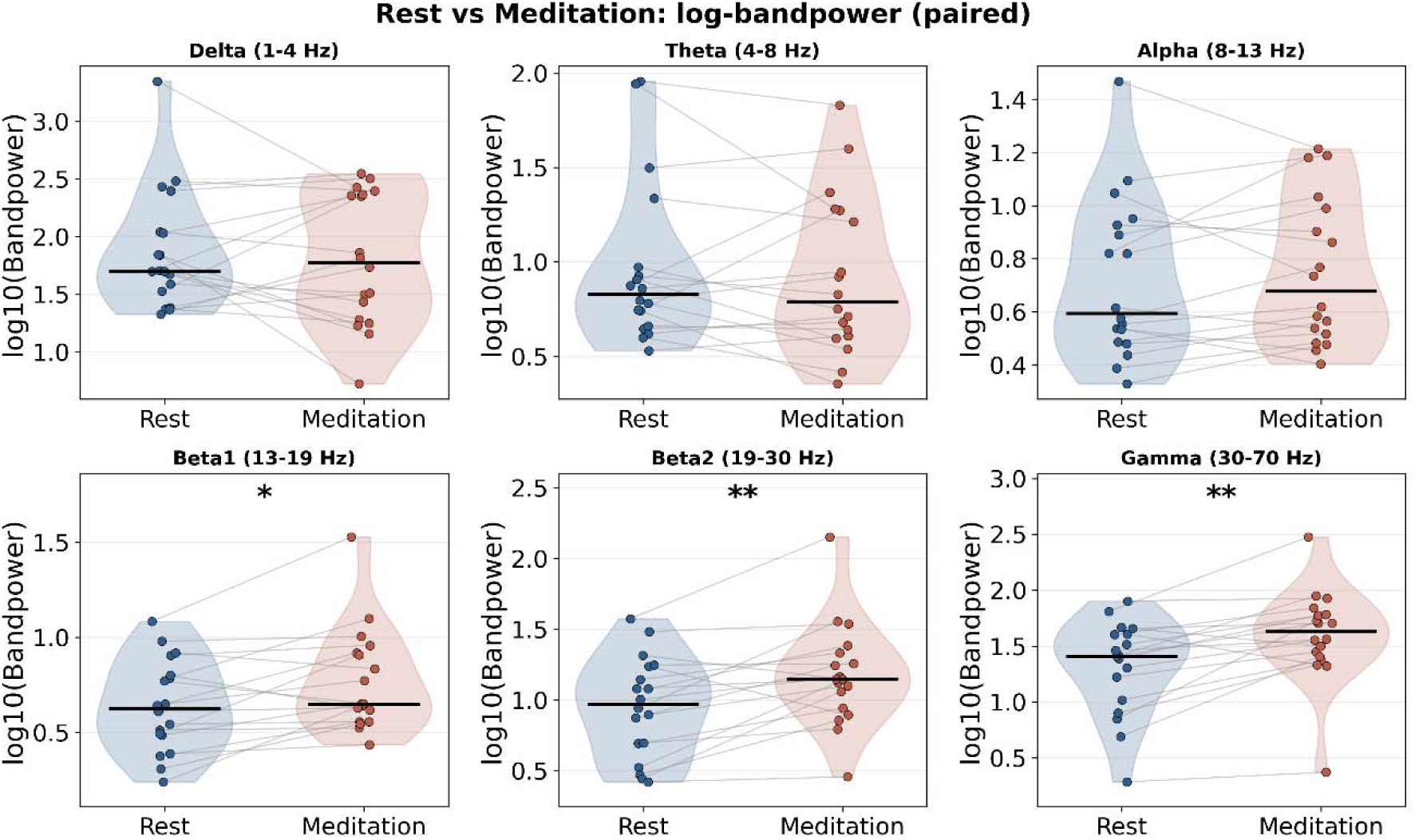
Spectral power during rest and meditation. Band power in six frequency bands (delta, theta, alpha, beta1, beta2, gamma) during the resting (blue) and meditation (red) conditions (N = 18). Violin plots show the distribution across participants, while filled circles show individual participants, joined by grey lines that connect each participant’s resting and meditation values. Horizontal black bars mark the group medians. Asterisks denote FDR-corrected significance (Wilcoxon signed-rank test, * FDR p < 0.05; ** FDR p < 0.01)..

### 4.2 Meditation reduces beta band temporal non-stationarity

Meditation also altered the temporal organization of EEG activity, reducing temporal non-stationarity, measured by means of the KLD, most clearly within the beta frequency range (Figure 3). After FDR correction, beta1 and beta2 KLD both decreased, indicating more temporally stable dynamics: beta1 fell by 0.012 (95% CI [-0.028, −0.003]; W = 21, r = −0.66, p = 0.003, FDR p = 0.020) and beta2 by 0.010 ([-0.019, 0.002]; W = 31, r = −0.56, p = 0.016, FDR p = 0.048). Gamma and alpha showed decreases of the same sign that were significant before correction but fell just short of it (gamma: r = −0.51, p = 0.030, FDR p = 0.051; alpha: r = −0.50, p = 0.034, FDR p = 0.051). Theta also decreased, but this was not significant (r = −0.46, p = 0.054, FDR p = 0.065), and delta was unchanged (r = −0.21, p = 0.39). Across the higher bands, the direction was consistent: meditation left the signal more stationary and less variable over time.

**Figure 3.**
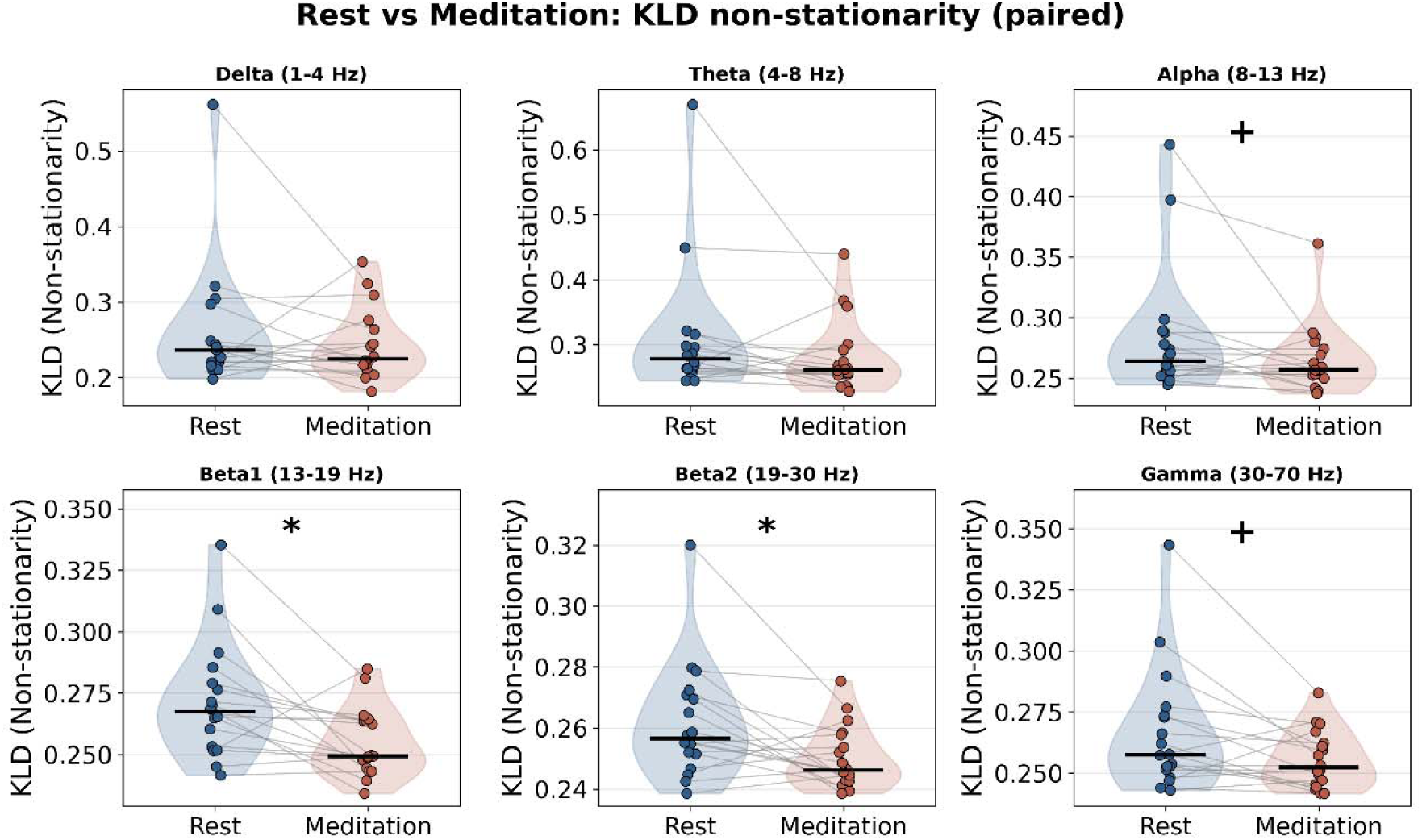
Temporal non-stationarity (KLD) during rest and meditation. Kullback–Leibler divergence, quantifying the temporal non-stationarity of the signal, in the same six frequency bands during the resting (blue) and meditation (red) conditions (N = 18). Violins show the across-participant distribution, while filled circles show individual participants joined by grey lines, and black bars mark the group median. Asterisks denote FDR-corrected significance (Wilcoxon signed-rank test, * FDR p < 0.05); plus signs (alpha and gamma) denote effects that were significant before correction but not after (+ p < 0.05, FDR p = 0.051).

### 4.3 Meditation shifts cortical dynamics toward an inhibition-dominant and lower intracortical gain regime

When the EEG spectra were translated into physiological parameters using the corticothalamic neural-field model, the meditation recordings showed a rebalancing of the cortical circuit toward stronger inhibition and lower effective intracortical gain (Figure 4). Specifically, after FDR correction across the five model parameters (G_ee_, G_ei_, G_ese_, G_esre_ and X), the cortical inhibitory gain became more negative during meditation than during rest (*G_ei_* decreased by 0.94; 95% CI [−3.49, −0.003]; W = 24, r = −0.63, p = 0.006, FDR p = 0.014) and the intracortical gain decreased (*X* fell by 0.060; [−0.115, −0.011]; W = 20, r = −0.67, p = 0.003, FDR p = 0.014). By contrast, the cortical excitatory gain (*G_ee_*) and the corticothalamic gain Gese did not change, and one corticothalamic loop gain (*G_esre_*) decreased only at trend level, i.e., non-significantly after correction (p = 0.054, FDR p = 0.090). In the corticothalamic framework, lower *X* indicates a shift toward reduced excitation relative to inhibition, making the overall pattern most consistent with a more inhibition-dominant cortical state during meditation, rather than a globally activated one. The remaining model parameters did not show any significant statistical differences.

**Figure 4.**
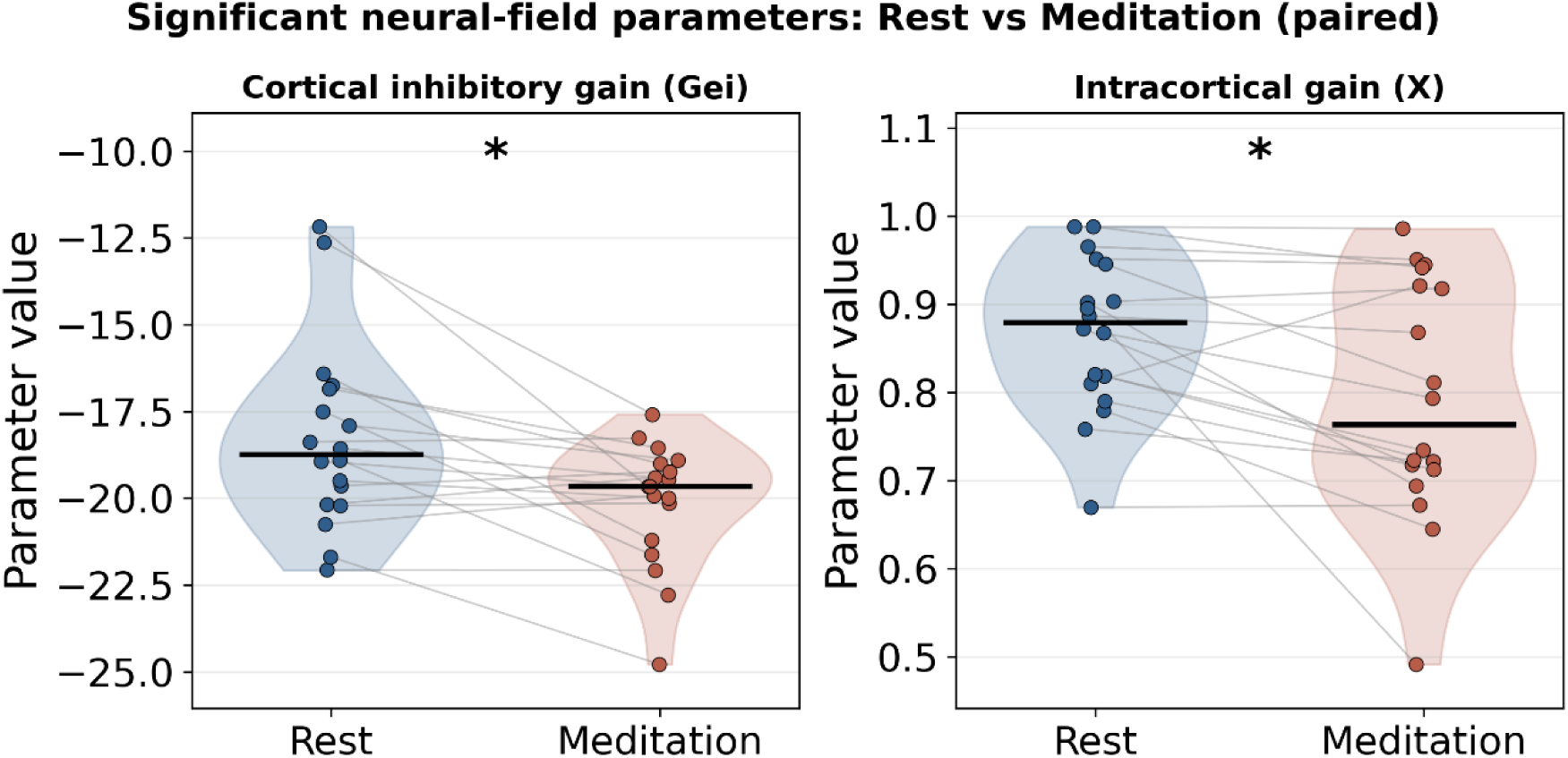
Corticothalamic model parameters during rest and meditation. The two neural-field model parameters that changed significantly with meditation: the cortical inhibitory gain ( ) and the intracortical gain ( ), during the resting (blue) and meditation (red) conditions (N = 18). Violin plots show the distribution across participants, while filled circles show individual participants, joined by grey lines that connect each participant’s resting and meditation values. Asterisks denote FDR-corrected significance (Wilcoxon signed-rank test, * FDR p < 0.05).

### 4.4 Meditation increases absorption and dissociation ratings

Both subjective ratings were higher during meditation than at rest. Median absorption rose from 6.0 at rest to 8.0 during meditation (median difference 1.25, 95% CI [1.0, 2.0]; W = 11.5, r = 0.73, p = 0.002), and median dissociation rose from 3.0 to 8.25 (median difference 4.5, [3.0, 6.5]; W = 0, r = 0.88, p < 0.001). Participants reported reaching both the rest and the meditative state quickly, with a median of 0.5 minutes in each condition (N = 17, as one participant did not report this).

### 4.5 The inhibitory and gain modulation scales with absorption

This physiological shift was also related to the depth of the subjective meditative state, indexed by absorption (Figure 5). During meditation, participants with a more negative *G_ei_* and a lower *X* reported greater absorption (*G_ei_*: rho = −0.51, 95% CI [-0.83, −0.01], p = 0.030; X: rho = −0.47, [-0.81, 0.01], p = 0.047). In exploratory analysis, the association was still clearer for rest-to-meditation change scores: participants who showed the largest increase in absorption also showed the largest increase in cortical inhibitory gain (*G_ei_* change vs absorption change: rho = −0.59, [-0.86, −0.19], p = 0.010), whereas the corresponding change-score correlation for *X* was weaker and not significant (rho = −0.28, p = 0.26). The remaining model parameters did not show any significant correlation with the depth of the meditative state.

**Figure 5.**
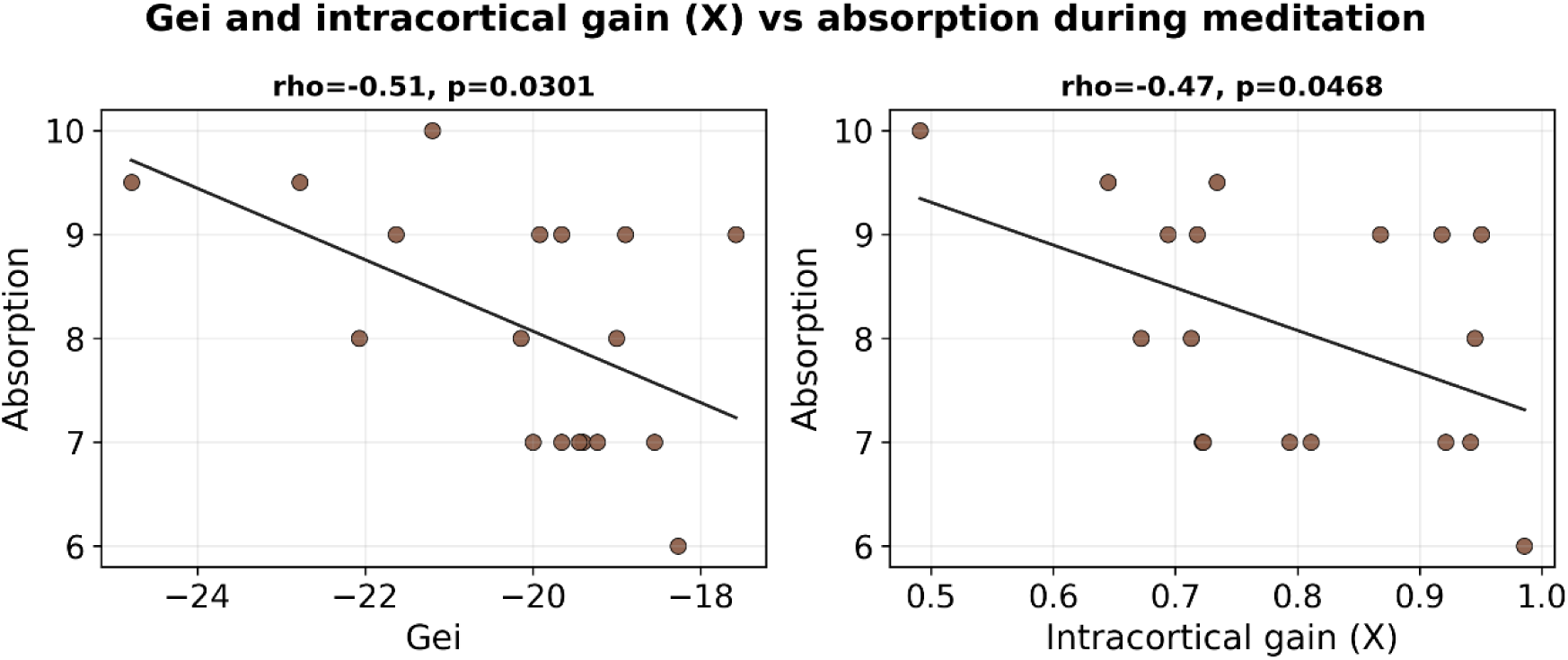
Association between corticothalamic model parameters and absorption during meditation. Scatter plots of the cortical inhibitory gain ( ) and the intracortical gain ( ), measured during meditation, against self-reported absorption during meditation (N = 18). Each point is one participant, and the solid line is the least-squares fit. Spearman rank-correlation coefficients (rho) and p-values are shown above each panel.

### 4.6 Dissociation shows the same direction at trend level

Dissociation pointed in the same direction, although the correlation was weaker (Figure 6). During meditation, both *G_ei_* and *X* were negatively associated with dissociation at trend level (*G_ei_*: rho = −0.45, 95% CI [-0.76, 0.02], p = 0.063; X: rho = −0.45, [-0.77, 0.03], p = 0.063), and the change-score correlations were weaker still (*G_ei_* change vs dissociation change: rho = −0.42, p = 0.080; *X* change: rho = −0.18, p = 0.48). Consistent with this overlap in direction, absorption and dissociation were themselves strongly correlated, both during meditation (rho = 0.67, 95% CI [0.25, 0.90], p = 0.002) and in their rest-to-meditation change (rho = 0.54, [0.15, 0.78], p = 0.019; Supplementary Figure S2). The two subjective measures thus behaved as closely related readings of a shared immersive dimension, with absorption of the more sensitive of the two to the model-derived inhibitory and gain modulation.

**Figure 6.**
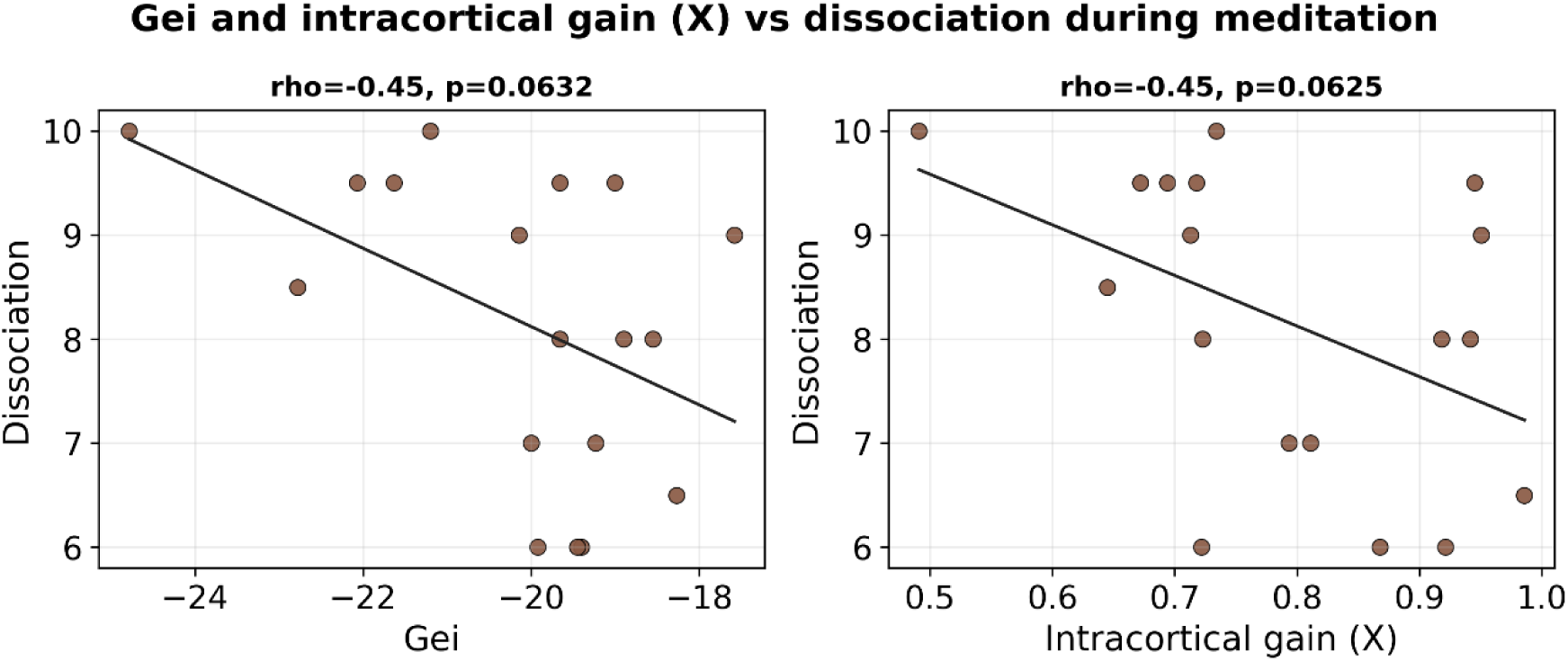
Association between corticothalamic model parameters and dissociation during meditation. Scatter plots of the cortical inhibitory gain ( ) and the intracortical gain ( ), measured during meditation, against self-reported dissociation during meditation (N = 18). Spearman rank-correlation coefficients (rho) and p-values are shown above each panel.

## 5. Discussion

In the present study, we asked how Rajyoga meditation reshapes ongoing cortical activity during an absorptive, self-detached meditative state. Using a single frontal EEG channel and three measures derived from the same cleaned data, we found that meditation increased higher-frequency power, with the effect growing from the beta1 through the gamma band and reduced the temporal non-stationarity of the signal in the beta range, leaving the activity more stable over time. Furthermore, when the spectra were interpreted using a corticothalamic neural-field model, meditation was associated with greater cortical inhibitory gain and lower effective intracortical gain. The magnitude of this inhibitory shift scaled with subjective absorptive depth. Together, these findings suggest that Rajyoga meditation involves several linked changes in cortical dynamics: fast activity is enhanced, its temporal expression becomes more stable, and the underlying cortical operating regime shifts toward inhibition-dominant gain control.

The increase in beta and gamma power fits the large existing literature. Elevated gamma is among the most reproducible electrophysiological markers of meditation, has been reported across several traditions, and tends to scale with the amount a person has practiced (Braboszcz et al., 2017; Lutz et al., 2004). In open-eye meditators, endogenous gamma is stronger than in matched controls and coexists with stimulus-driven gamma, which suggests that the two are generated by partly separate mechanisms (Biswas et al., 2026). Shorter interventions and concentrative practice can also move beta- and gamma-range power (Maher et al., 2025; Stapleton et al., 2020; Tang et al., 2007). Our data extends this picture in two ways: the effect appeared in a minimal, single-channel frontal montage, and it was graded: strengthening steadily with frequency. Because gamma accompanies sustained attention, which in advanced meditators often becomes effortless, the most natural reading is that the higher-frequency increase indexes the focused absorption that Rajyoga cultivates (Fries, 2005; Jensen et al., 2007b; Lutz et al., 2008).

Power alone, however, misses a second property of the meditative signal. Alongside the increase in beta and gamma power, meditation reduced the beta-band non-stationarity of the EEG, meaning that the distribution of spectral energy became more even across time. This is in line with studies that have examined the temporal dynamics and complexity of the EEG during meditation (Escrichs et al., 2019; Irrmischer et al., 2018; Young et al., 2021). Note that KLD measures non-stationarity and is not itself a measure of complexity. Experienced Zen practitioners show altered EEG complexity during practice (Huang & Lo, 2009), and skilled Rajyoga meditators can shift their ongoing dynamics rapidly and voluntarily (Nair et al., 2017). A signal that is both higher in beta and gamma power and more stationary in the beta range points to a state that is not just more activated at higher frequencies but also more steadily maintained, a pattern that may reflect the steady attention that Rajyoga practice aims to build. That the power and non-stationarity effects converged on the same bands, from the same epochs, strengthens this reading.

The neural-field model let us move past a purely descriptive towards an estimated physiological account of the EEG changes. Fitting the corticothalamic model (Abeysuriya & Robinson, 2016; Robinson et al., 1997, 2001, 2002) to each condition, meditation produced a more negative cortical inhibitory gain (*G_ei_*) and a lower intracortical gain (*X*), with the two moving together because *X* is a normalized function of the cortical gains. Because G_ee_ did not change, the lower X mainly reflects the stronger inhibition. We still report X because it summarises, in one number, how strong excitation in the cortex is offset by inhibition, which allows comparison with earlier work using this model (Abeysuriya & Robinson, 2016; Robinson et al., 2002). It might seem strange for a state to show both increased gamma power and stronger inhibition, since inhibition is often taken as a marker of reduced activity. But gamma oscillations are generated precisely by the interaction between excitatory and inhibitory neuronal populations, with inhibitory interneurons setting their rhythm (Buzsáki, 2006; Buzsáki & Wang, 2012; Cardin et al., 2009; Sohal et al., 2009). A cortex that is more strongly governed by inhibition is therefore expected to produce more, not less, gamma activity. This followsthe interpretation that Biswas et al. (2026) reached from an independent open-eye meditation dataset, proposing that stronger inhibition underlies the enhanced gamma activity observed in meditators. Independent neurochemical evidence is consistent with a meditation-related increase in cortical inhibition: meditation and yoga practice have been linked to elevated GABAergic cortical inhibition and brain GABA levels (Guglietti et al., 2013; Streeter et al., 2010).

We also reported correlataions between the physiological estimations from the model and the subjective accounts from the practitioners. Within our pre-specified tests, the model’s inhibitory and gain modulation tracked subjective absorption: participants with a more negative *G_ei_* and a lower intracortical gain *X* during meditation reported deeper absorption, and those whose absorption increased most from rest to meditation showed the largest inhibitory change. This convergence echoes prior work linking absorptive depth to changes in the temporal structure of cortical activity during focused-attention meditation (Irrmischer et al., 2018). The relationship with dissociation ran in the same direction but was weaker and did not reach significance. We believe, however, that this difference between the two measures may have a simple explanation: absorption and dissociation were themselves strongly correlated, both during meditation and in their change scores, which is consistent with the long-standing view that absorption and dissociative immersion are closely related dimensions of altered consciousness (Bowins, 2012; Mohr, 2018; Soffer-Dudek et al., 2015; Tellegen & Atkinson, 1974). Absorption appears to be the more sensitive of the two to the cortical modulation we observed, perhaps because it captures the depth of attentional engagement most directly (Tellegen & Atkinson, 1974), whereas dissociation adds a component of self-detachment that is less tightly coupled to the model parameters.

The present study has several limitations First, a period of rest always preceded meditation, so the two conditions differ not only in mental state but also in time on task. Part of the change could reflect adaptation, drowsiness, or the passage of time rather than meditation as such, and a counterbalanced or separately cued design would be needed to distinguish these. Second, the recordings were derived from a single frontal channel, Fp1-Fp2. This makes the approach practical and portable, but it cannot localize the effects, and much of the meditation gamma literature reports parieto-occipital rather than frontal sources (Biswas et al., 2026; Braboszcz et al., 2017). Third, due to the aforementioned limitation of single-channel recordings, independent component analysis could not be applied, so we cannot fully exclude a muscle contribution to the high-frequency findings. However, the recordings were inspected and epochs were rejected to mitigate this to the best of our abilities. It is worth noting that the effects were different across bands, was accompanied by a model change, and survived spike and flatline screening, but the gamma band is the most influenced by EMG noise and as such, its results should be interpreted with caution. Fourth, the sample was modest and drawn from long-term practitioners of a single tradition, without a non-meditating control group, so the findings reflect within-person state changes rather than to trait differences or to meditation in general. Fifth, absorption and dissociation were measured with simple single-item ratings given after each condition, not with a validated questionnaire. Sixth, we did not include an active task condition, which would be a useful comparison for an eyes-open practice such as Rajyoga. Finally, KLD describes non-stationarity only; future work could add complexity measures such as Lempel-Ziv complexity and permutation entropy, which have been useful for separating states of consciousness (Sitt et al., 2014).

Our findings showed that Rajyoga meditation, a practice involving a dissociative state with focused attention, modulates brain dynamics towards high-frequency activity and enhances intracortical inhibition. Long-term Rajyoga practitioners entered a meditative state marked by stronger beta and gamma activity, more stationary beta dynamics and a cortical regime with stronger inhibitory gain and lower intracortical gain. The inhibitory shift scaled with absorptive depth, and dissociation followed the same trend at the trend level, suggesting that absorbed attention and self-detachment draw on a shared cortical mechanism. Inhibitory gain control is a possible mechanism for this shared effect, because it can keep an internal focus stable while reducing the influence of bodily and external signals on experience. In conclusion, we believe that meditative self-detachment is thus best understood as a controlled, inhibition-dominant state of the cortex.

## Acknowledgements

We are grateful to all the participants who took part in this study and contributed their time and effort.

## Conflict of Interest

The authors declare no conflict of interest.

## Funding

This study was supported by the Canada Excellence Research Chair (CERC).

## Data Availability

The data supporting the findings of this study will be made available upon reasonable request.

## Notes

### Competing Interest Statement

The authors have declared no competing interest.

